# A simple and rapid CRISPR-Cas12a-based detection test for diastatic *Saccharomyces cerevisiae*

**DOI:** 10.1101/2022.11.23.517627

**Authors:** Ida Uotila, Kristoffer Krogerus

## Abstract

Diastatic *Saccharomyces cerevisiae* is a common contaminant in the brewing industry. Currently available detection methods are either time-coansuming or require specialized equipment. The aim of this study was to develop a new rapid and simple assay for the detection of diastatic yeast from beer and yeast samples. More specifically, we aimed to develop a simple and rapid assay that requires minimal laboratory equipment or training, and ideally yields results as accurate as PCR-based methods. The developed assay consisted of three main steps: DNA extraction, pre-amplification of DNA, and CRISPR-Cas12a-based detection and visualisation. We compared different preamplification and visualisation techniques, and the final assay involved a one-pot reaction where LAMP and Cas12a were consecutively used to pre-amplify and detect a fragment from the *STA1* gene in a single tube. These reactions only required a heat block, a pipette, and a centrifuge. The assay result was then visualised on a lateral flow strip. We used the developed assay to monitor an intentionally contaminated beer fermentation, and it was shown to yield results as accurate as PCR using previously published primers. Furthermore, the assay yielded results in approx. 75 minutes starting from a beer sample. The developed assay therefore offers reliable and rapid quality control for breweries of all sizes and can be performed without any expensive laboratory equipment. We believe the assay will be particularly useful for smaller breweries that don’t already have well-equipped laboratories and are looking to implement better quality control.

## 1. Introduction

Diastatic *Saccharomyces cerevisiae* (var. *diastaticus*) is one of the most common spoilage microbes in the brewing industry (Meier-Dörnberg et al. 2017; Powell and Kerruish 2017; Meier-Dörnberg et al. 2018; Krogerus and Gibson 2020). These strains, which are genetically closely related to brewing strains of *S. cerevisiae*, produce an extracellular glucoamylase enzyme that enables the hydrolysis of otherwise unfermentable dextrin to fermentable glucose. Contamination by diastatic yeast can therefore lead to extended fermentations and organoleptic faults. In serious cases, increased carbon dioxide formation in packaged beers can lead to gushing and exploding packages that may injure the consumer. Contamination by diastatic *S. cerevisiae* appears to be most common during bottling, and smaller breweries, with less quality control and more experimentation with different yeast strains, tend to be more susceptible to contamination (Meier-Dörnberg et al. 2017). To maintain high beer quality and prevent economic and reputational losses, it is therefore vital that diastatic *S. cerevisiae* can be rapidly and reliably detected within the brewery.

Various detection methods for diastatic yeast are currently available, and they include traditional growth-based methods and modern molecular methods (Powell and Kerruish 2017; Krogerus et al. 2019; Krogerus and Gibson 2020; Traynor et al. 2021; Burns et al. 2021). However, as diastatic strains are genetically and physiologically similar to brewing strains, it can be difficult to distinguish contaminants from production strains. The growth-based methods, which rely on culturing samples on selective media, are popular because of their simplicity and low cost, but they are time-consuming and prone to false positives depending on the selective media. The molecular methods typically rely on the detection of the *STA1* gene, which encodes the extracellular glucoamylase responsible for the diastatic phenotype. Detection can be accomplished with polymerase chain reaction (PCR), either endpoint or quantitative, and several commercial kits are available on the market. A typical workflow includes extraction of DNA from a (potentially pre-enriched) yeast sample, performing PCR using primer pair for *STA1* (e.g. SD-5A/SD-6B from Yamauchi et al. (1998)), and separation and visualisation of PCR products using gel electrophoresis and DNA staining. The advantages of the molecular methods are speed and accuracy, as results can be obtained within hours instead of days. However, these methods also require specialized equipment and training, and therefore might not be accessible to smaller breweries.

Recently, a number of nucleic acid detection techniques exploiting various CRISPR-Cas nucleases have been described (Gootenberg et al. 2017; Chen et al. 2018; Li et al. 2018; Kellner et al. 2019). The CRISPR-Cas nucleases can be guided to cleave specific nucleic acid sequences using a single guide RNA molecule. The detection techniques exploit a feature of the Cas12a and Cas13 enzymes, where the activated nuclease exhibits non-specific *trans* cleavage activity on any nearby single-stranded DNA (ssDNA) or RNA, respectively. For Cas12a, this means ssDNA is cut non-specifically when the nuclease is bound to a double-stranded DNA activator with complementary base-pairing to the guide crRNA (Chen et al. 2018). This *trans*-activity can be read out, e.g. as an increase in fluorescence, if a ssDNA reporter containing a fluorophore and a quencher is used. A further increase in sensitivity can be obtained if the target DNA is pre-amplified prior to the CRISPR-Cas reaction, e.g. using PCR or isothermal methods (Li et al. 2018; Wang et al. 2019; Kellner et al. 2019). These systems have been used for the detection of various human viral diseases, including human papillomavirus and SARS-CoV-2 (Chen et al. 2018; Broughton et al. 2020; Ali et al. 2020), and food spoilers, such as *Salmonella* sp. and *Escherichia coli* (Wang et al. 2020; Ma et al. 2021; Liu et al. 2022). Recently, a Cas12a-based assay was also developed for the detection of common beer spoilage lactic acid bacteria (Meng et al. 2021).

Here, our objective was to develop a CRISPR-Cas12a-based assay for the detection of diastatic yeast from beer and yeast samples. More specifically, we aimed to develop a simple and rapid assay that requires minimal laboratory equipment or training, and ideally yields results as accurate as PCR-based methods and within approx. one hour starting from yeast sample. The developed assay consisted of three main steps: DNA extraction, pre-amplification of DNA, and CRISPR-Cas12a-based detection and visualisation. We compared different pre-amplification techniques, including PCR and two isothermal techniques (recombinase polymerase amplification (RPA) and loop-mediated isothermal amplification (LAMP)), and two different visualisation techniques (fluorescence and lateral flow strips). The final assay involved a one-pot reaction where LAMP and Cas12a were consecutively used to pre-amplify and detect a fragment from the *STA1* gene in a single tube. The result was then visualised on a lateral flow strip. The assay was used to monitor an intentionally contaminated beer fermentation, and was shown to yield results as accurate as PCR using previously published primers. Furthermore, the assay yielded results in approx. 75 minutes starting from a beer sample. The developed assay therefore offers reliable and rapid quality control for breweries of all sizes and can be performed without any expensive laboratory equipment.

## 2. Materials and methods

### 2.1 Yeast strains

A list of strains used in this study can be found in Table 1. These included both ale and diastatic yeast strains. Yeasts were pre-cultured in 25 mL YPD media (1% yeast extract, 2% peptone and 2 glucose) at 25□°C and 120 rpm overnight inside an Infors HT Minitron incubator (Infors HT, Bottmingen, Switzerland).

**Table 1.** Yeasts used in this study.

| Commercial name | Short code | *STAI* expression | Species | Source |
| --- | --- | --- | --- | --- |
| VTT-A75060 | A60 | - | *Saccharomyces cerevisiae* | VTT Culture Collection, Espoo, Finland |
| WLP023 Burton Ale Yeast | WLP023 | - | *Saccharomyces cerevisiae* | White Labs Inc., San Diego, CA, USA |
| 3711 French Saison | WY3711 | + | *Saccharomyces cerevisiae* | Wyeast Laboratories, Odell, OR, USA |
| TUM PI BA 109 | TPB109 | + | *Saccharomyces cerevisiae* | BLQ Weihenstephan, Freising, Germany |

For testing of the developed protocols, a dilution series of A60 and WY3711 cultures was prepared based on optical density (absorbance at 600 nm). Pre-cultures of both strains were diluted to equal optical density with sterile deionized water, after which ten-fold dilutions of the WY3711 suspension in the A60 suspension were carried out. Step-wise dilution was carried out to obtain 10^−1^ to 10^−6^ fractions of WY3711 in A60.

### 2.2 DNA extraction

DNA was extracted from yeast cultivations with ‘GC Prep’ (Blount et al. 2016) using 425-600 μm acid-washed glass beads (Sigma-Aldrich, USA) and a 5% suspension of Chelex^®^ 100 sodium form as particle size 50-100 mesh (dry) (Sigma-Aldrich, USA) diluted in sterile Milli-Q water. Yeast was were separated from samples by centrifugation (1 min, 5000 × *g*). 50 μL 5% Chelex 100 solution and approximately 50 μL glass beads were added to yeast pellet. The yeast sample was vortex-mixed for 4 minutes, after which the mixture was incubated for 2 minutes at 98 °C. The genomic DNA was dissolved in the liquid phase and was collected from the supernatant after centrifugation (1 min, 15000 × *g*). DNA concentration and A260/280 ratio of samples was measured with a Qubit 2.0 fluorometer and Nanodrop spectrophotometer, respectively.

### 2.3 Design of crRNA

Targeting with the Cas12a endonuclease occurs with a guide RNA (gRNA) molecule complementary to the target site, and also requires a 5’ TTTV protospacer adjacent motif (PAM). As *STA1* is a chimeric gene of *FLO11* and *SGA1*, here, a suitable protospacer sequence specifically targeting *STA1*, and not *FLO11* nor *SGA1*, needed to be designed. Potential protospacer sequences were obtained using CCTop (Stemmer et al. 2015), with the *STA1* promoter and open reading frame (ORF) sequence of *S. cerevisiae* WY3711 (obtained from Krogerus et al. 2019) being the target, and the *S. cerevisiae* S288C genome (Engel et al. 2014) being used for detection of off-target sites. Any potential protospacer sequences, lacking predicted off-target activity (i.e. more than four mismatches) in *FLO11, SGA1* or any other genomic location, were then further queried for matches in the genomes of the 157 *S. cerevisiae* strains sequenced by (Gallone et al. 2016) using ‘blastn’ in the online suite of NCBI-BLAST. 100% matches were only allowed in the 21 strains carrying *STA1* (Krogerus et al. 2019). A single protospacer sequence, (TTTC)AATTAGAACCACAACATGAC, was ultimately selected for the trials (Table 2). The sequence is located 40 bp upstream of the *STA1* start codon. A crRNA molecule containing the selected protospacer sequence was ordered from IDT (Integrated DNA Technologies, Inc., Coralville, Iowa, USA) (Table 2).

**Table 2.**
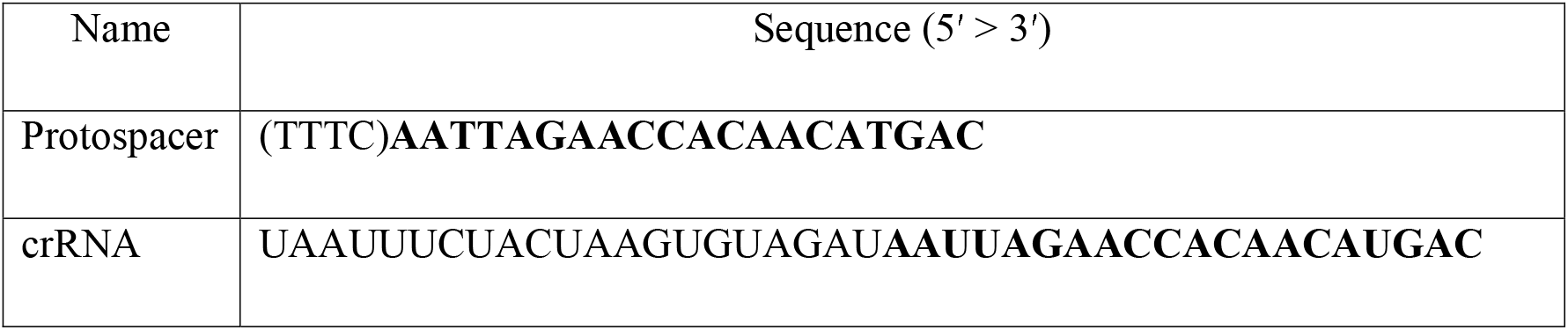
Protospacer sequence and crRNA used in this study.

### 2.4 Design of primers

PCR and RPA primers (Table 3 and Supplementary Table 1) were designed using Primer-BLAST (Ye et al. 2012) with default settings. The aim was to produce amplicons containing the above Cas12a protospacer sequence, with amplicon lengths around 200 bp. The primers were designed to be around 20 and 30 bp long for PCR and RPA reactions, respectively.

**Table 3.**
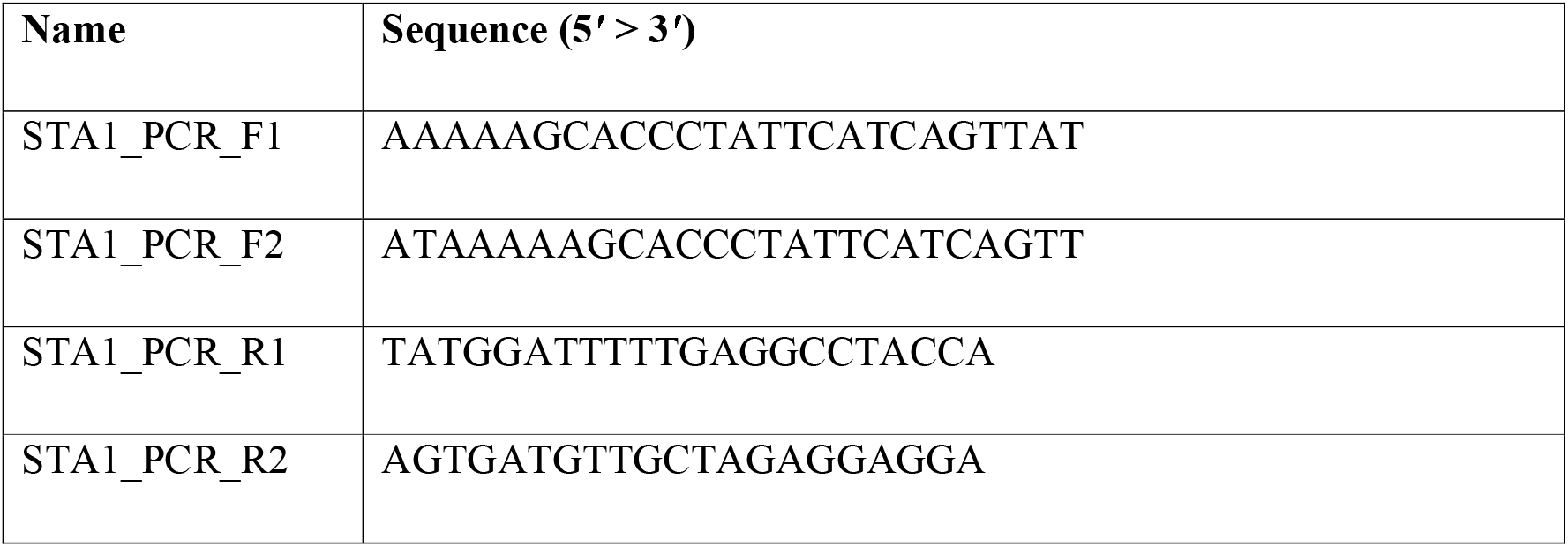
Primers designed to PCR reactions.

LAMP primers were designed using both GLAPD (Jia et al. 2019) and the NEB LAMP Primer Design online tool version 1.2.0 (https://lamp.neb.com/). Again, the amplicons were designed to contain the Cas12a protospacer sequence. In GLAPD, the *S. cerevisiae* S288C genome was used to test the specificity of the generated primers. Two different LAMP primer sets were obtained and tested (Table 4).

**Table 4.**
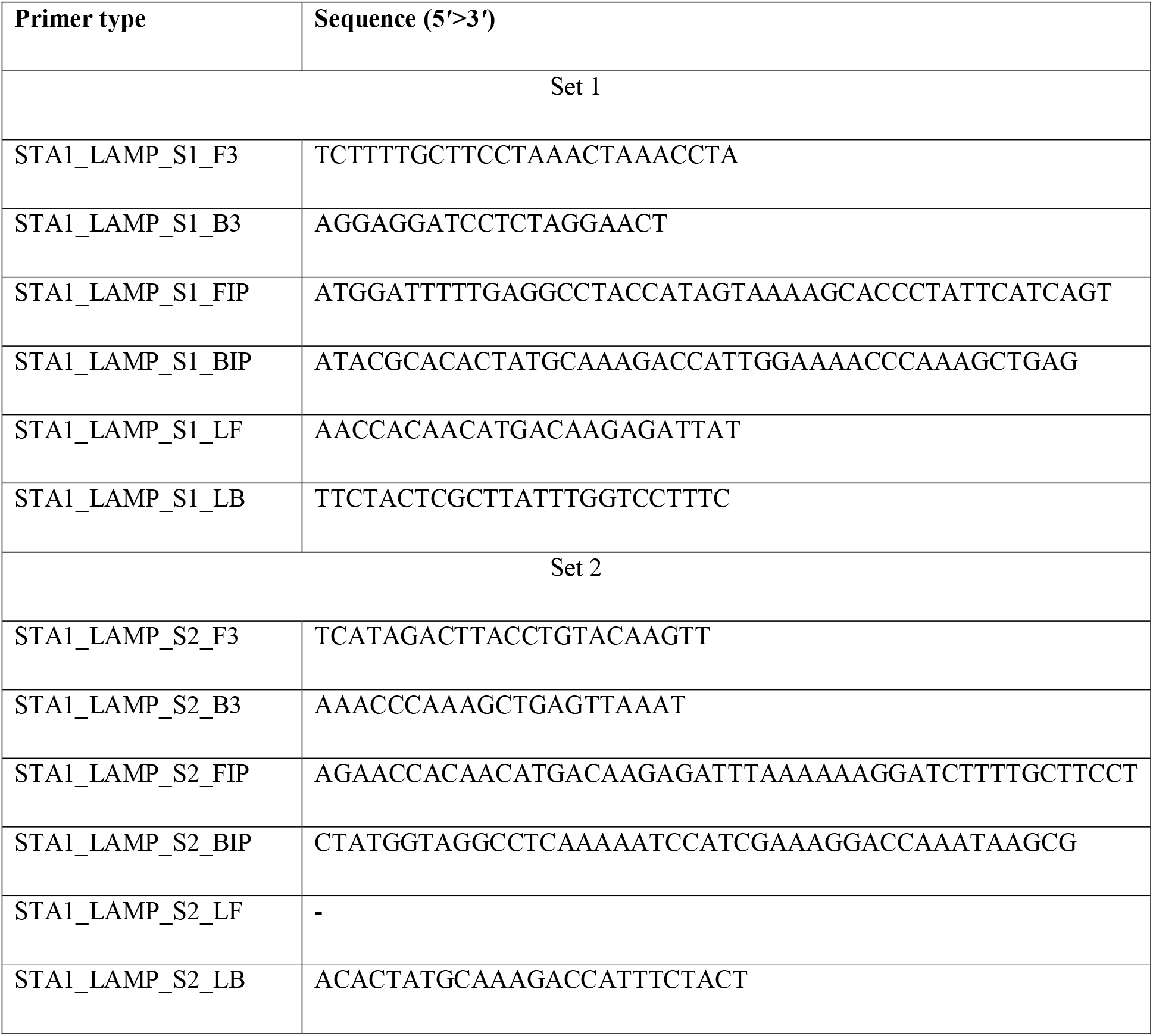
Primer set 1 and 2 designed for LAMP reaction.

All primers were purchased from Thermo Scientific (Finland).

### 2.5 PCR amplification

PCR reactions were carried out with Phusion polymerase. Both Phusion High-Fidelity PCR Master Mix with HF Buffer (Thermo Scientific, Vantaa, Finland) and Phusion™ Plus Green PCR Master Mix (Thermo Scientific, Vantaa, Finland) were used. The reactions were carried out according to manufacturer’s instructions, and reaction volumes were adjusted to 14 μL with primer concentrations of 0.5 μM. PCR reactions were performed on a Mastercycler^®^ Nexus (Eppendorf, Hamburg, Germany).

The PCR reactions with the newly designed primers were optimized with increasing annealing temperature from 62.1 °C to 64 °C, decreasing elongation time from 30 sec to 6 sec and increasing cycles from 30 to 35. As a control, *STA1* was also detected with previously the previously published primer pair SD-5A 5’-CAACTACGACTTCTGTCATA-3’ and SD-6B 5’-GATGGTGACGCAATCACGA-3’ (Yamauchi et al. 1998) and the PCR programme used with the SD-5A/6B primers was: 98 °C for 2 min, 30 cycles of (98 °C for 10 s, 57.5 °C for 30 s, 72 °C for 30 s), 72 °C for 10 min and cooling to 8 °C. PCR products were separated on 1.5% agarose gels in 0.5× TBE buffer with gel electrophoresis.

### 2.6 RPA amplification

Pre-amplification of the *STA1* gene was attempted with recombinase polymerase amplification (RPA) (Piepenburg et al. 2006). RPA is an isothermal amplification method, and amplification is therefore carried out at a constant temperature. The RPA reactions were carried out with a commercial TwistAmp® Liquid Basic Kit (TALQBAS01, TwistDx, Maidenhead, UK) according to the manufacturer’s instructions with minor modifications to volumes. Reaction volumes were adjusted to 15 μL containing 7.5 μL of 2x Reaction Buffer, 1.8 mM of dNTPs, 1.5 μL 10x Basic E-mix, 0.48 μM of each primer, 0.75 μL 20x Core Reaction Mix, 0.96 μL of 280 mM MgOAc and 1.5 μL of DNA template. Reaction conditions were optimized for RPA with incubation temperatures of 37 °C, 39 °C and 42 °C. Amplicons were cleaned with AMPure XP beads (Beckman Coulter Inc., Brea, CA, USA) prior to separation on 1.5% agarose gels in 0.5× TBE buffer with gel electrophoresis.

### 2.7 LAMP amplification

Pre-amplification of the *STA1* gene was also attempted with loop-mediated isothermal amplification (LAMP) (Notomi 2000). Like RPA, LAMP is also an isothermal amplification method. LAMP reactions were set up using the NEB WarmStart^®^ LAMP Kit (DNA & RNA) (E1700S, New England Biolabs, Ipswich, Massachusetts, USA) and using the two designed primer sets. Reactions were carried out in 10 μL reaction volume containing 5 μL WarmStart LAMP 2X Master Mix, 1.6 μM FIP, 1.6 μM BIP, 0.2 μM F3, 0.2 μM B3, 0.4 μM LF, 0.4 μM LB and 1 μL of DNA template. Primer set 2 lacked the LF primer and was replaced with H_2_O. Reaction conditions were optimized by testing different incubation temperatures (65 °C, 67 °C and 69 °C) and dimethyl sulfoxide (DMSO) concentration (0%, 5% and 7.5%) (Wang et al. 2015). The incubation time was 30 minutes according to the kit protocol.

### 2.8 CRISPR-Cas12a-based nucleic acid detection

CRISPR-Cas12a reactions were carried out in 20 μL reaction volumes with 2 μL NEBuffer™ r2.1 (New England Biolabs, Ipswich, Massachusetts, USA), 500 nM crRNA, 500 nM Cas12a enzyme, varying ssDNA oligonucleotide concentrations, 0.5 μL RNase inhibitor (TaKaRa Bio Inc., Japan) and 1 μL sample. Two different Cas12a enzymes were used: EnGen® Lba Cas12a (Cpf1) (New England Biolabs, Ipswich, Massachusetts, USA) and Alt-R™ L.b. Cas12a (Cpf1) Ultra (Integrated DNA Technologies, Inc., Coralville, Iowa, USA). Reactions were carried out in RNase free tubes with RNase- and DNase-free pipette tips to avoid any RNase or DNase contamination. Different ssDNA oligonucleotide reporters were used depending on desired read-out (described below in 2.8.1 and 2.8.2).

#### 2.8.1 Fluorescent read-out

A fluorophore quencher (F.Q.) -labeled ssDNA oligonucleotide reporter was used in order to detect Cas12a activity through fluorescence (Chen et al. 2018). The reporter used here was HPLC-purified 5’ / 6-FAM / TTATT / Iowa Black® FQ / 3’ purchased from IDT (Integrated DNA Technologies, Inc., Coralville, Iowa, USA). CRISPR-Cas12a reactions were prepared on Microfluor 2Black “U” Bottom Microtiter Plates (Thermo Scientific, Rochester NY, USA). The reactions contained 500 nM of F.Q.-labeled reporter, and they were incubated at 37 °C for 30 minutes during which the change in fluorescence was monitored. The intensity of fluorescence was measured with excitation at 495 nm and emission at 520 nm with a VarioSkan (Thermo Scientific) using SkanIt™ Software version 2.3.4.

#### 2.8.2 Lateral flow read-out

A biotin-labelled ssDNA oligonucleotide reporter was used to detect Cas12a activity with lateral flow strips (Broughton et al. 2020). The reporter was HPLC-purified 5’ / 6-FAM / TTATT / Biotin / 3’ purchased from IDT. Different reporter concentrations were tested, and later reactions were carried out with a concentration of 400 nM. CRISPR-Cas12a reactions were incubated at 37 °C for 30 minutes and the assay result was read out with lateral flow strips from a HybriDetect - Universal Lateral Flow Assay Kit (MGHD 1, Milenia Biotec GmbH, Gießen, Germany). 5 μL of CRISPR-Cas12a reaction product was added to the application area of lateral flow dipstick and strips were incubated for 1 min in 100 μL of HybriDetect assay buffer in an upright position and results were observed immediately and photographed with a phone camera.

### 2.9 One-Pot LAMP-Cas12a reaction

A one-pot reaction, combining the LAMP and CRISPR-Cas12a reactions in a single vessel to reduce operating steps and cross-contamination risks, was designed inspired by previous studies (Wang et al. 2019; Wang et al. 2021). As the LAMP reaction is incubated at 65 °C, a temperature at which the EnGen® Lba Cas12a enzyme is inactivated, the LAMP reaction had to be physically separated from the Cas12a reaction mix with mineral oil and an air phase. A 10 μL LAMP reaction was set up in the bottom of a 1.5 mL tube and sealed with 25 μL of mineral oil, while 20 μL CRISPR-Cas12a reaction mix was assembled in the screw cap. Reagent concentrations of CRISPR-Cas12a reaction mix were increased 1.5-fold to taken into account the increased total volume. Reactions were performed with both the F.Q. reporter and biotin-labelled reporter. Reaction tubes were first incubated for 30 minutes at 65 °C, during which the LAMP reaction occurs, after which tubes were cooled for 1 minute at room temperature followed by mixing with five full inversions. The liquid was briefly spun down, after which tubes were incubated at 37 °C for 30 minutes. Results were read depending on the reporter used in the reaction. When the F.Q. reporter was used, fluorescence was observed under UV light. When the biotin-labelled reporter was used, lateral flow read-out was carried out as previously above (5 uL of sample was taken under the oil phase).

### 2.10 Wort fermentations

Small-scale wort fermentations were carried out to test the developed method on wort and beer samples. The aim was to carry out fermentations inoculated with diastatic yeast (TPB109), non-diastatic yeast (WLP023) and finally with non-diastatic yeast contaminated with diastatic yeast (WLP023 and TPB109). Overnight pre-cultures of the yeasts were prepared in 50 mL YPM media (1% yeast extract, 2% peptone and 2% maltose) at 25 °C and 120 rpm shaking. Pre-cultured yeasts were pelleted and resuspended in deionized H_2_O to 20% (w/v) slurries. The viability and cell concentration of the slurries was measured with a NucleoCounter® YC-100™ (Chemometec A/S, Allerod, Denmark). Wort was prepared from Spraymalt Extra Light (Muntos, Suffolk, UK) extract to a concentration of 12 °P and wort was boiled with 10 g/L Cascade pellets for 30 minutes. Wort was filtered to clarify, and pasteurized at 80 °C for 30 minutes. Yeast was inoculated at a rate of 1.2 × 10^7^ cells/mL into 80 mL of wort. The intentionally contaminated fermentations were additionally spiked with 8.4 × 10^3^ cells of TBP109 to obtain a contamination level of 120 cells/mL. Fermentations were carried out in duplicate 100 mL sterile Schott bottles capped with glycerol-filled airlocks at 20 °C for 11 days.

1 mL samples were drawn regularly during the fermentations. Prior to sampling, the bottles were carefully agitated, and 1 mL of wort was pipetted into an Eppendorf tube and centrifuged (3 min, 4000 × *g*). Supernatants were removed and analyzed for fermentable sugars as described below. The yeast pellets were washed twice, by resuspending yeast pellet in 1 mL of sterile Milli-Q water and removing the water slurry by centrifugation (3 min, 4000 × *g*). Yeast pellets were stored at -20°C until DNA extraction. DNA was extracted with ‘GC Prep’, and the genomic DNA was used to test the developed LAMP-Cas12a protocol.

The fermentable sugar and ethanol concentrations in samples were analyzed with high-performance liquid chromatography (HPLC) using a Waters Alliance® HPLC system that consisted of a Waters 2695 Separation Module, Waters System Interphase Module and Waters 2414 differential refractometer (Waters Co., Milford, MA, USA). A Bio-Rad Aminex HPX-87H HPLC column for organic acid analysis (300 × 7.8 mm; Bio-Rad Hercules, CA, USA) was equilibrated with 5 mM H_2_SO_4_ (Titrisol, Merck, Germany) in water at ambient temperature 55 °C and analyzed samples were eluted with 5 mM H_2_SO_4_ in water at flow rate of 0.3 mL/min.

## 3. Results and discussion

Here, the aim was to develop a simple and rapid assay for detection of diastatic *S. cerevisiae*, and the final overall workflow is shown in Figure 1. The developed assay consisted of three main steps: DNA extraction, pre-amplification of DNA, and CRISPR-Cas12a-based detection and visualisation. The final assay involved a one-pot reaction where LAMP and Cas12a were consecutively used to pre-amplify and detect a fragment from the *STA1* gene in a single tube. The assay result was then visualised on a lateral flow strip.

**Figure 1.**
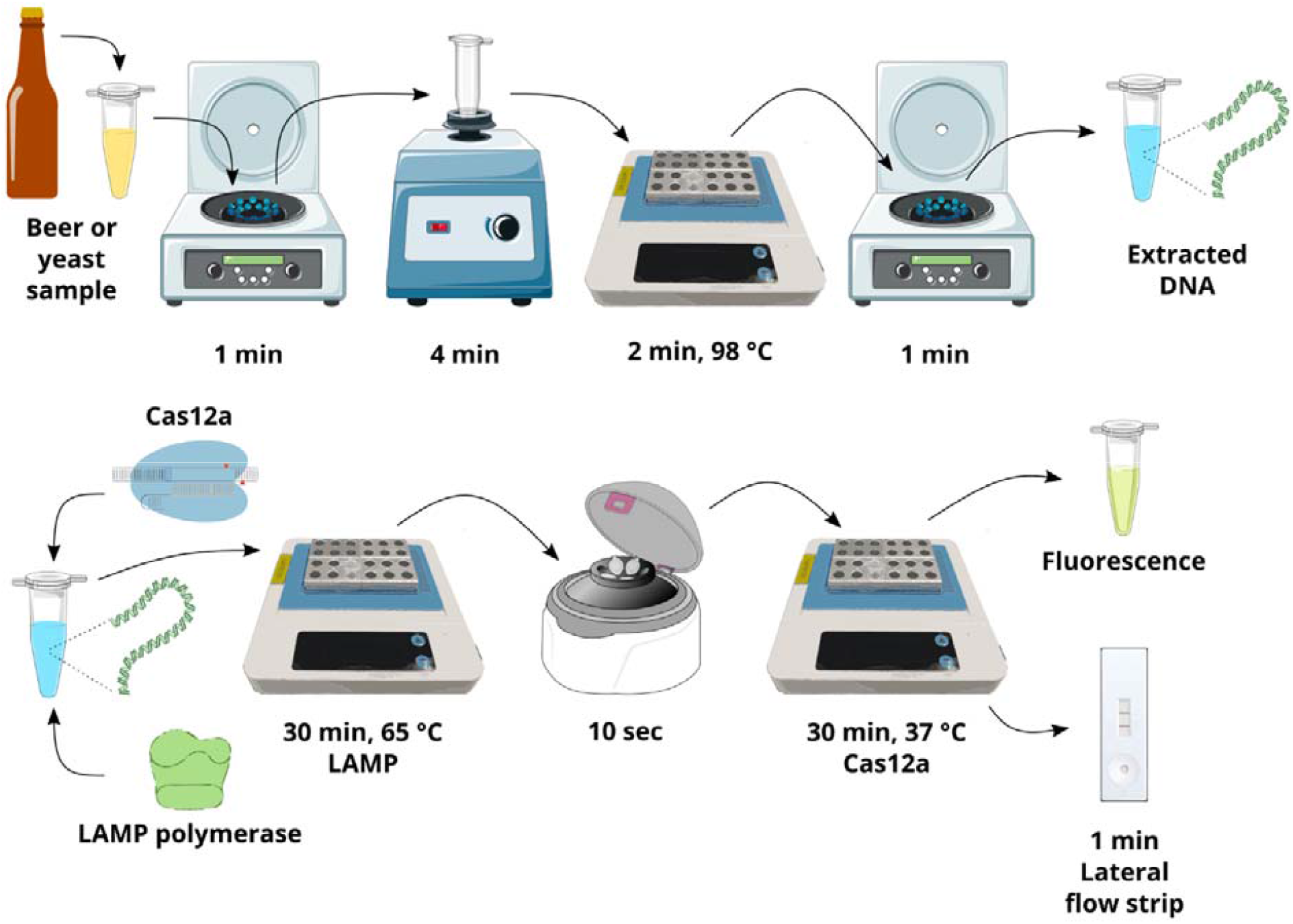
An overview of the developed detection assay for *STA1* from beer or yeast samples.

### 3.1 DNA extraction and testing of pre-amplification strategies

The first step of the developed assay involved DNA extraction from samples. ‘GC Prep’ was selected for its speed, simplicity and resulting DNA quality (Blount et al. 2016). DNA was successfully extracted from the included brewing yeast strains using the ‘GC Prep’ method. DNA concentrations ranged from 2.6 to 6.8 ng/μL, while A260/280 ratios varied from 2.1 to 2.4. The total time to process a sample was approximately 10 minutes. The extracted DNA was then used to compare different pre-amplification strategies.

#### 3.1.1 Pre-amplification by PCR

Four different primer combinations (F1-R1, F1-R2, F2-R1 and F2-R2) were designed to target the *STA1* gene of diastatic *S. cerevisiae* and produce approximately 200 bp amplicons containing the selected Cas12a protospacer sequence with PCR. Initial trials used a PCR programme of: 98°C for 2 min, 30 cycles of (98°C 10 sec, 62.1°C 20 sec and 72°C for 30 sec), and 72°C for 10 minutes. With this programme, all four primer combinations produced an amplicon with the positive control (WY3711) (Figure 2A). However, the negative control (A60) also produced target amplicons for two of the pairs (F1-R1 and F2-R1), and larger unwanted byproducts (∼1000 and 1200 bp) with the other two primer pairs. To remove these unwanted byproducts, the PCR programme was optimized for the F2-R2 primer pair. By increasing the annealing temperature from 62.1 °C to 64 °C, decreasing elongation time from 30 sec to 6 sec and increasing amplification cycles from 30 to 35, formation of the unwanted byproduct in the negative control was eliminated (Figure 2B). The modifications also decreased the overall run time of the PCR programme. Two different Phusion master mixes, Phusion High-Fidelity PCR Master Mix with HF Buffer and Phusion Plus Green PCR Master Mix, were compared and these yielded identical results. Pre-amplification with PCR using primer pair F2-R2 was therefore successful, however the step would require a thermal cycler and approximately 45-60 minutes of run time. Furthermore, using PCR as a pre-amplification method would yield few advantages compared to just using PCR and gel electrophoresis directly for *STA1* detection.

**Figure 2.**
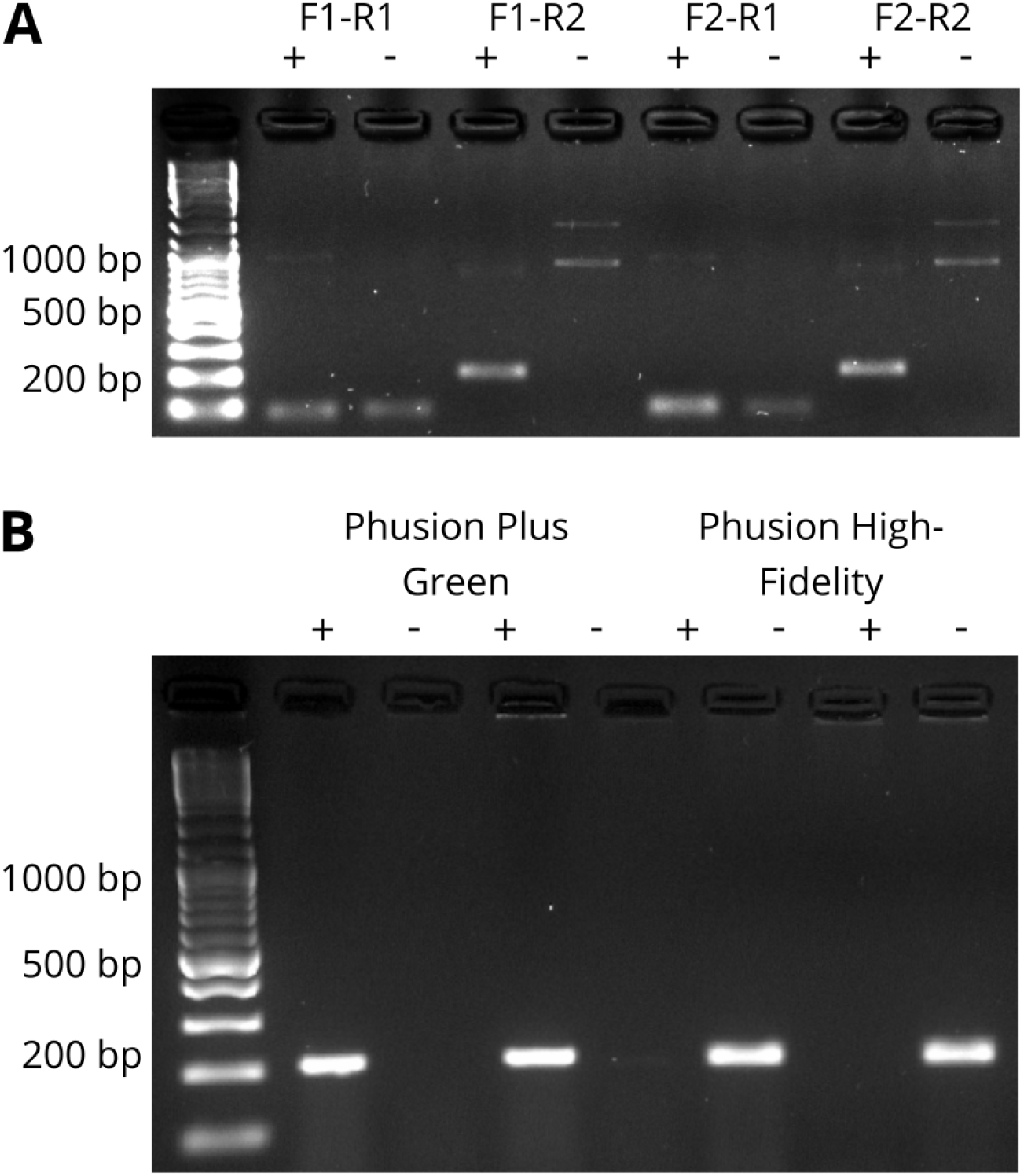
(**A**) PCR reactions with four different primer pairs targeting *STA1* using DNA from the *STA+* control *S. cerevisiae* WY3711 and *STA-* control *S. cerevisiae* A60. PCR reactions were carried out with Phusion High-Fidelity PCR Master Mix with HF Buffer. (**B**) Optimized PCR reactions with the F2/R2 primer pair using DNA from the same strains as in (**A**). The first four reactions were carried out with Phusion™ Plus Green PCR Master Mix and the last four were carried out with Phusion High-Fidelity PCR Master Mix with HF Buffer.

#### 3.1.2 Isothermal pre-amplification by RPA and LAMP

To simplify and speed up the pre-amplification step, two different isothermal DNA amplification methods were explored: recombinase polymerase amplification (RPA) and loop-mediated isothermal amplification (LAMP). For RPA, a total of twelve primer combinations were tested on a *STA+* (WY3711) and *STA-* strain (A60) with a commercial TwistAmp^®^ Liquid Basic Kit. The four most promising primer combinations (i.e. producing the most and least product for the positive and negative sample, respectively) were optimized by testing different incubation temperatures between 37 °C and 42 °C.

Overall, RPA was not able to differentiate the positive and negative samples as clearly as PCR (Figure 3). Poor amplification was observed with all primer pairs at 42 °C. For primer pairs F3-R3 and F3-R4, amplicons of equal intensity were produced in both samples at all temperatures. With primer pairs F1-R3 and F2-R2, a clearer difference between positive and negative samples was observed, and the most intense bands were produced with an incubation temperature of 39 °C. However, both primer combinations still produced faints bands with the negative control. An incubation temperature of 40 °C was further tested, but negative samples still produced faint bands (Figure 3B). The F2-R2 primer pair was chose for further experiments.

**Figure 3.**
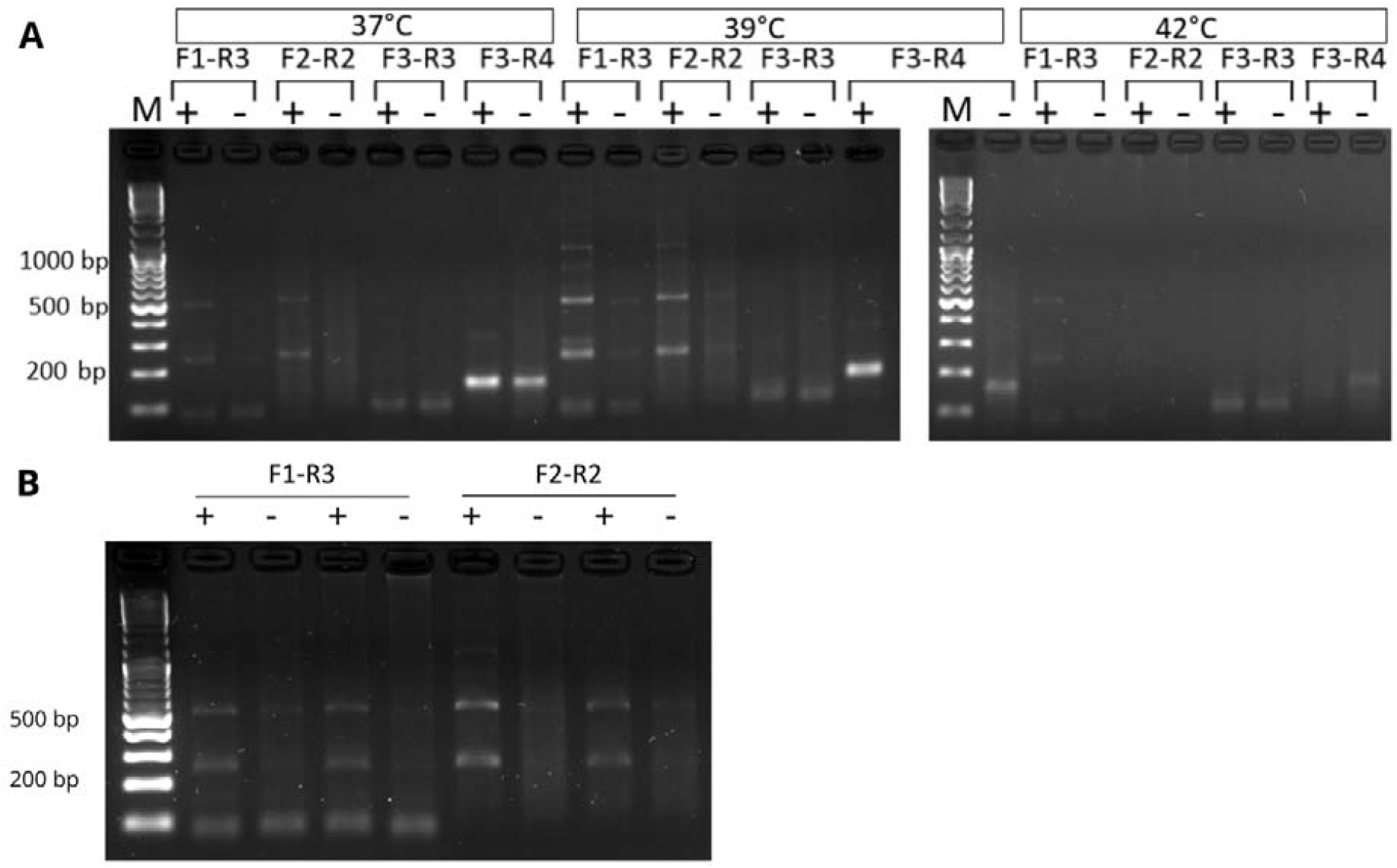
Optimization of the RPA reaction using DNA from the *STA1+* control *S. cerevisiae* WY3711 and *STA-* control *S. cerevisiae* A60. (**A**) Optimization is performed by changing the incubation temperatures between 37 °C and 42 °C using four primer pairs. (**B**) RPA reaction at 40°C with the two most promising primer pairs.

LAMP reactions are carried out with four to six primers, and here two different designed primer sets were tested on a *STA+* (WY3711) and *STA-* strain (A60) with a commercial NEB WarmStart^®^ LAMP Kit. An incubation temperature of 65 °C and DMSO content of 0% was used in the first trials. Set 1 produced an amplified product only in the positive control, while set 2 did not result any amplification in either sample (Figure 4). The amplified product, which is formed from repeats of the desired target sequence, can be seen as bright bands of different lengths. Different incubation temperatures (65, 67, and 69 °C) and DMSO contents (0, 5, and 7.5%) were further tested, however no further improvement in specificity and amplicon intensity was obtained compared to the initial trial (Supplementary Figure 1). Primer set 1, an incubation temperature of 65 °C, and DMSO content of 0% was selected for further experiments.

**Figure 4.**
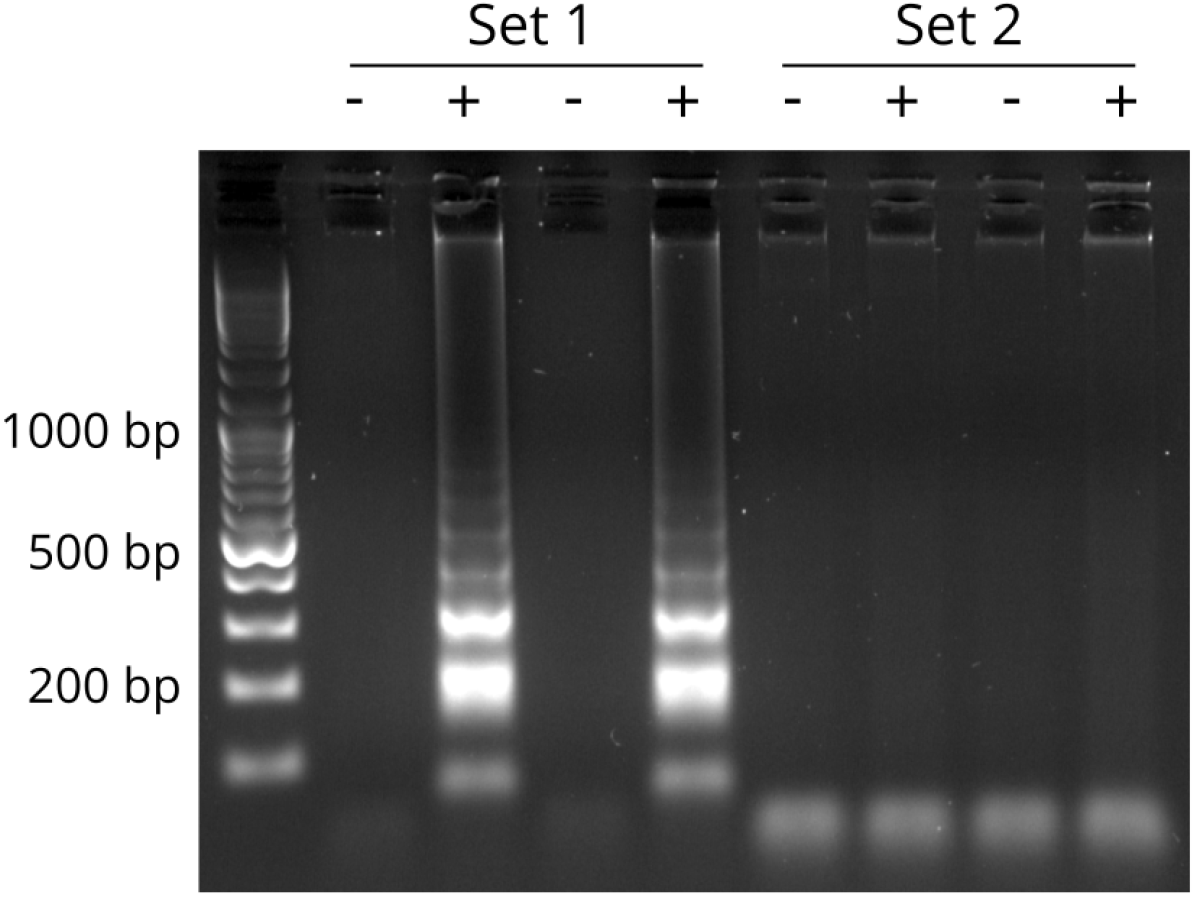
LAMP reactions using DNA from the *STA1-* control *S. cerevisiae* A60 and *STA1+* control *S. cerevisiae* WY3711 with primer set 1 and 2.

#### 3.1.3 Discussion of pre-amplification results

Of the three compared pre-amplification methods, LAMP appeared to be the most promising in that it could clearly differentiate a *STA1+* from a *STA1-* strain, and the reaction could be performed isothermally. PCR and LAMP primers for *STA1* gene detection have been previously developed (Yamauchi et al. 1998; Hayashi et al. 2009), but the existing primers could not be used here as they do not produce amplicons containing the Cas12a protospacer sequence. Primer design to produce amplicons containing the Cas12a protospacer sequence proved challenging, which is evident from the poor specificity observed during the RPA amplifications and some of the PCR primers, and the failed amplification for the second set of LAMP primers. The main challenge in primer design was that the *STA1* gene is chimeric, consisting of fragments from both *FLO11* and *SGA1* (Yamashita et al. 1987; Krogerus and Gibson 2020). The protospacer sequence is located just upstream of *STA1*, and the region is homologous to that upstream of *FLO11*. Hence, to develop primers specific for *STA1*, off-target activity against *FLO11* had to be checked.

### 3.2 CRISPR-Cas12a assay

#### 3.2.1 Fluorescent read out

The first CRISPR-Cas12a reactions were carried out on DNA samples either with or without pre-amplification using PCR or RPA. The DNA samples were obtained from a ten-fold dilution series of *STA1+* WY3711 in *STA-* A60. A non-template control, where the template DNA was replaced with water, was also included. A reporter molecule containing a fluorophore and quencher was used in the reaction so that Cas12a activity could be monitored by measuring increase in fluorescence. Reactions were incubated for 30 minutes, and fluorescence intensity was measured after incubating 3 and 30 minutes (Table 5). Results indicated that pre-amplification of DNA samples was necessary, as no Cas12a activity was observed in any of the non-pre-amplified samples. Differences between the positive and negative controls were, on the other hand, observed based on increasing fluorescence when pre-amplification with both PCR or RPA was used. PCR-amplified samples showed a clearer increase in fluorescence compared to RPA-amplified samples, with the latter producing only modest fluorescence in the diluted samples after 30 minutes of incubation. Hence, RPA pre-amplification was considered unsuitable for reliable detection using the CRISPR-Cas12a assay.

**Table 5.**
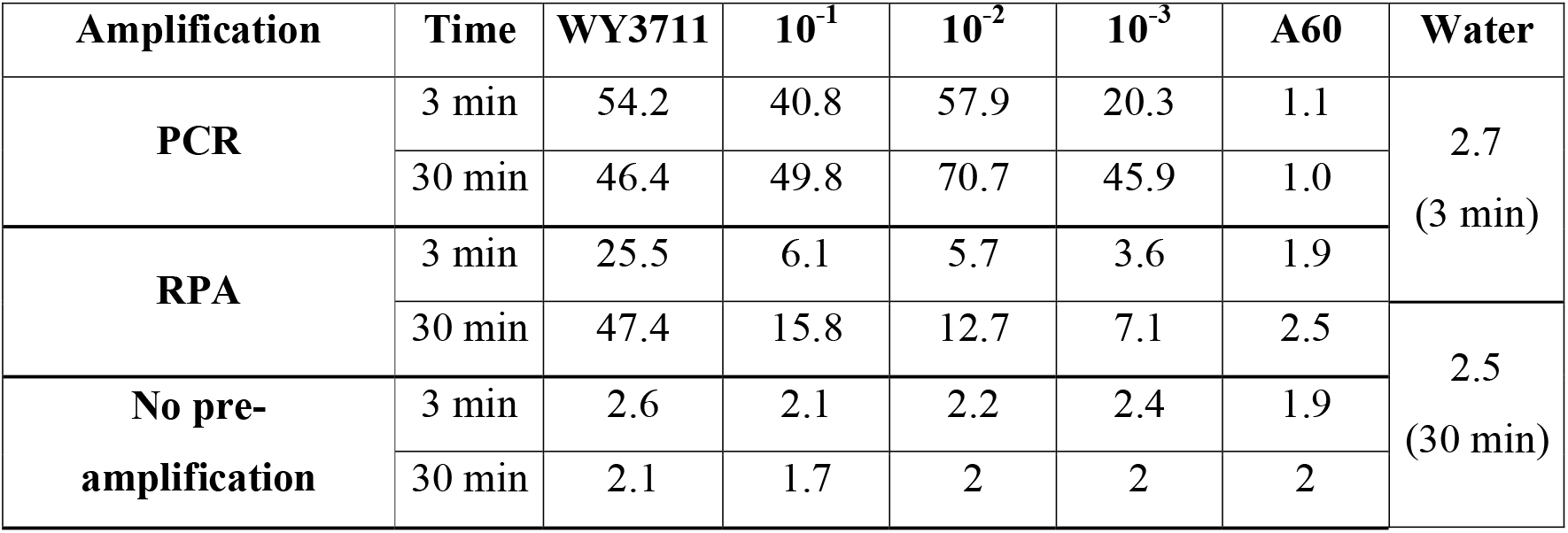
Fluorescence intensity (RFU) after Cas12a reactions using DNA pre-amplified with PCR or RPA from the dilution series of *STA1-*positive WY3711 in the *STA1*-negative A60.

As RPA was not a suitable isothermal pre-amplification method, we next tested CRISPR-Cas12a reactions on samples pre-amplified with LAMP. DNA was extracted from an extended dilution series (WY3711, 10^−1^-10^−6^, A60), and the genomic DNA was pre-amplified with both LAMP and PCR. DNA samples without pre-amplification (positive control, 10^−1^ dilution and negative control) and a non-template control were again included. Fluorescence intensity was measured after 1 and 10 minutes of incubation with the Cas12a enzyme (Table 6). In samples pre-amplified with LAMP, an increase in fluorescence intensity was observed for WY3711, 10^−1^, 10^−2^, 10^−3^ and 10^−4^ samples. At 10 minutes, these same samples pre-amplified with PCR yielded lower fluorescence intensity. LAMP therefore appeared to be the most sensitive out of the tested pre-amplification methods, and also has the benefit of being an isothermal method.

**Table 6.**
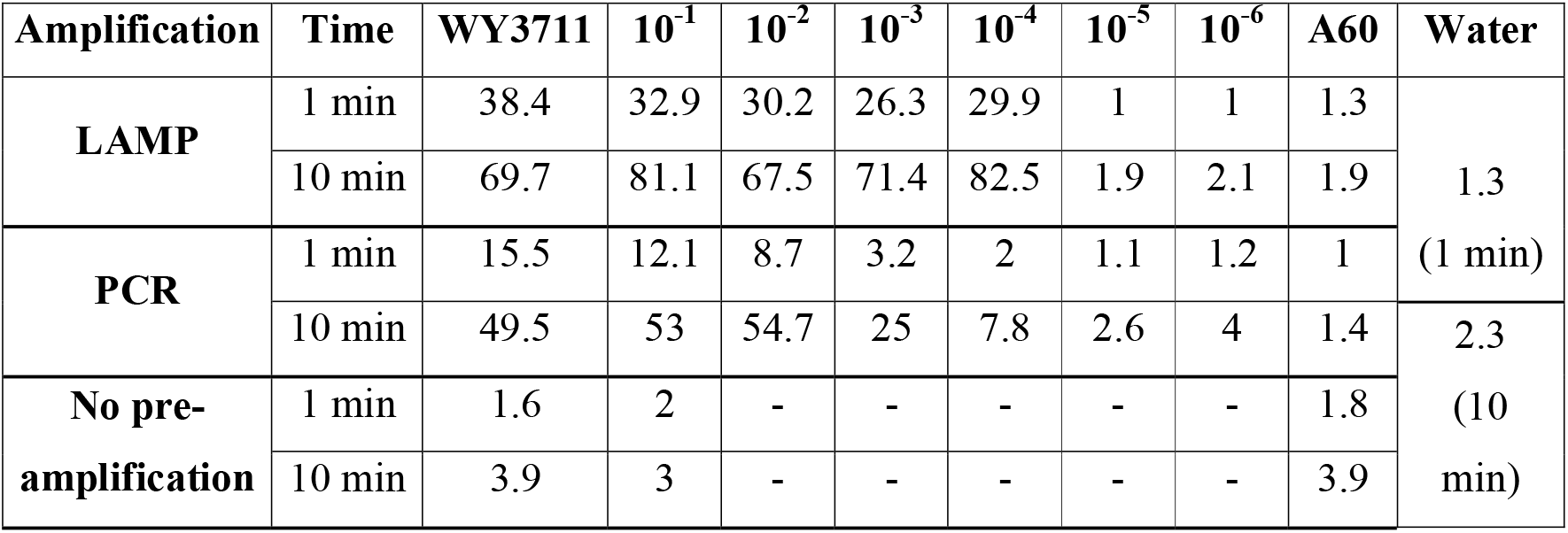
Fluorescence intensity (RFU) after Cas12a reactions using DNA pre-amplified with LAMP or PCR from the dilution series of *STA1-*positive WY3711 in the *STA1*-negative A60.

As pre-amplification with LAMP appeared the most promising, the dynamics of fluorescence increase during the Cas12a reactions was studied in more detail using DNA extracted from the dilution series (WY3711, 10^−1^ to 10^−6^ and A60) and replicate reactions. Four replicate reactions for each DNA sample were incubated with Cas12a for 30 minutes at 37 °C, and fluorescence intensity was continuously monitored (Figure 5). A non-template control was again included, but results are excluded from the plot as measured fluorescence was identical to the negative control (A60). Increase in fluorescence (i.e. Cas12a activity) was rapid, with fluorescence plateauing after around five minutes in WY3711, 10^−1^, and 10^−2^ samples. An increase in fluorescence was also observed with the 10^−3^, 10^−4^, and 10^−5^ samples, but it occurred slower and with a longer lag time. Fluorescence did not increase in 10^−6^ and the negative control A60 during the 30 minute incubation.

**Figure 5.**
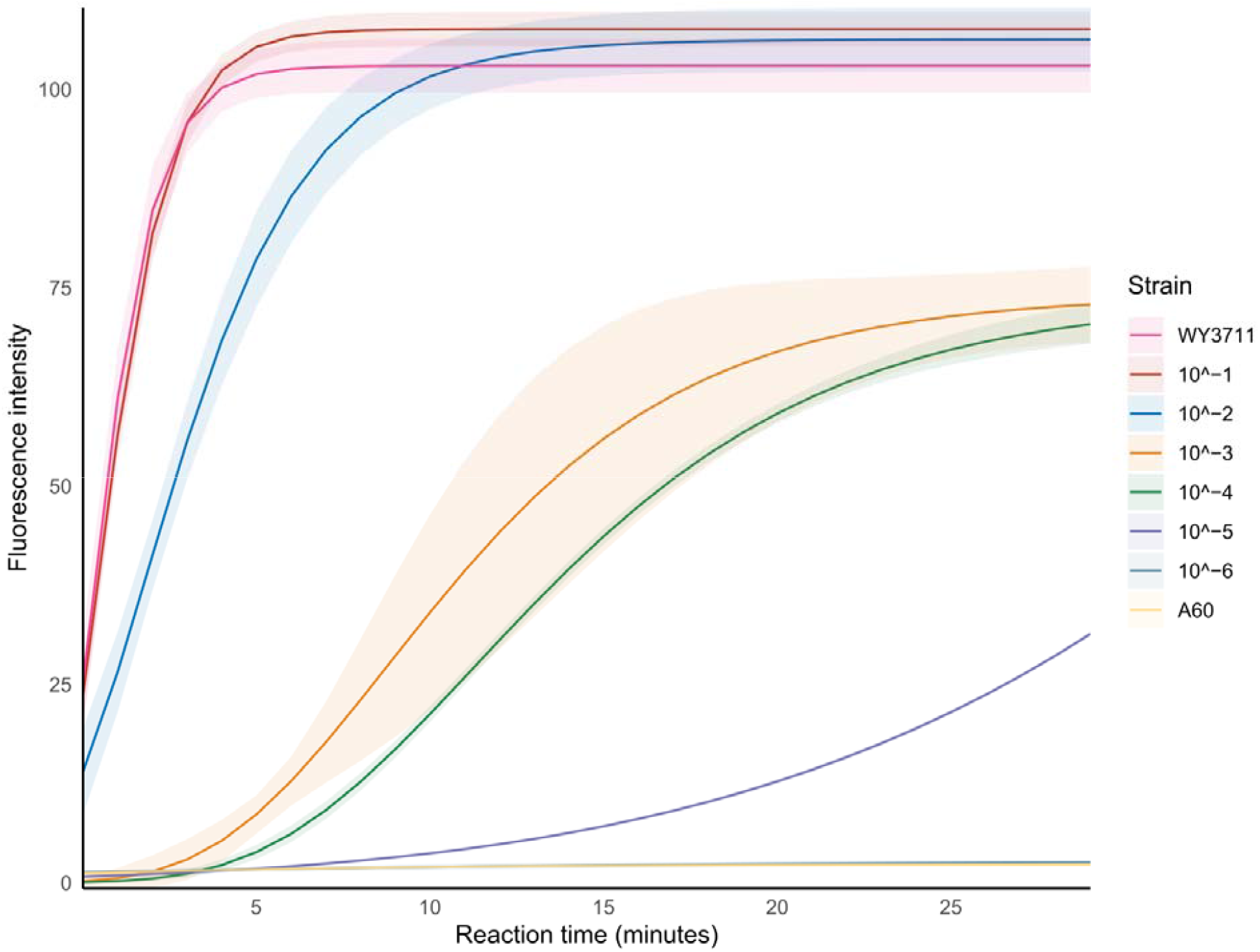
Change in fluorescence during Cas12a reactions using DNA pre-amplified with LAMP from the dilution series of *STA1+ S. cerevisiae* WY3711 in the *STA1-S. cerevisiae* A60. Values are replicates of four independent reactions, and the standard deviation is represented as the lighter shaded area.

#### 3.2.2 Lateral flow read out

Following successful demonstration of *STA1* detection using Cas12a, we next explored whether assay results could be visualized with lateral flow strips instead of fluorescence for a simpler and more user-friendly assay. A HybriDetect Universal Lateral Flow Assay Kit was used, and prior to testing visualization of Cas12a activity with lateral flow strips, the concentration of the biotin-labelled ssDNA reporter was optimized with aqueous solutions of the reporter. This was done according to the manufacturer’s instructions, as a non-optimal reporter concentration can result in so called ‘hook effect’, where all of the reporter is not retained at the control and a test line is unintentionally visible. 5 μL of the reporter dilutions were directly added to dipsticks and strips were incubated for 1 minute in manufacturer’s assay buffer. The objective was to minimize the intensity of the test line, while retaining a visible control line (C). The T line remained visible at all tested reporter concentrations (Supplementary Figure 2). While it was not possible to minimize the T line completely, the best results were obtained with a reporter concentration of 200-500 nM. Hence, for subsequent experiments, a concentration of 400 nM was used. It was further tested whether supplementing 5% polyethyleneglycol (PEG) to the buffer would decrease the strength of the T line, but no positive effect could be observed.

Since the T line could not be completely minimized in an aqueous solution of just the reporter, we next tested whether Cas12a-treated positive and negative samples could even be distinguished with the lateral flow strips. CRISPR-Cas12a reactions with the biotin-labelled reporter were performed on LAMP pre-amplified samples of DNA extracted from the previously used dilution series. The CRISPR-Cas12a reactions were incubated at 37 °C for 30 minutes and the results were immediately read with lateral flow strips (Figure 6A). Clear differences between positive and negative controls could be observed even though T line could not be eliminated with optimization. On the strips loaded with Cas12a-treated samples from the positive control and dilutions 10^−1^ to 10^−5^, the T line was clearly more intense than the C line, which had disappeared completely. LAMP amplicons were also separated with gel electrophoresis to compare result, and these same samples showed clear amplification during LAMP (Figure 6B). Therefore, based on our results, a positive result with the lateral flow strips (i.e. *STA1* detection) is obtained when the T line is more intense than the C line.

**Figure 6.**
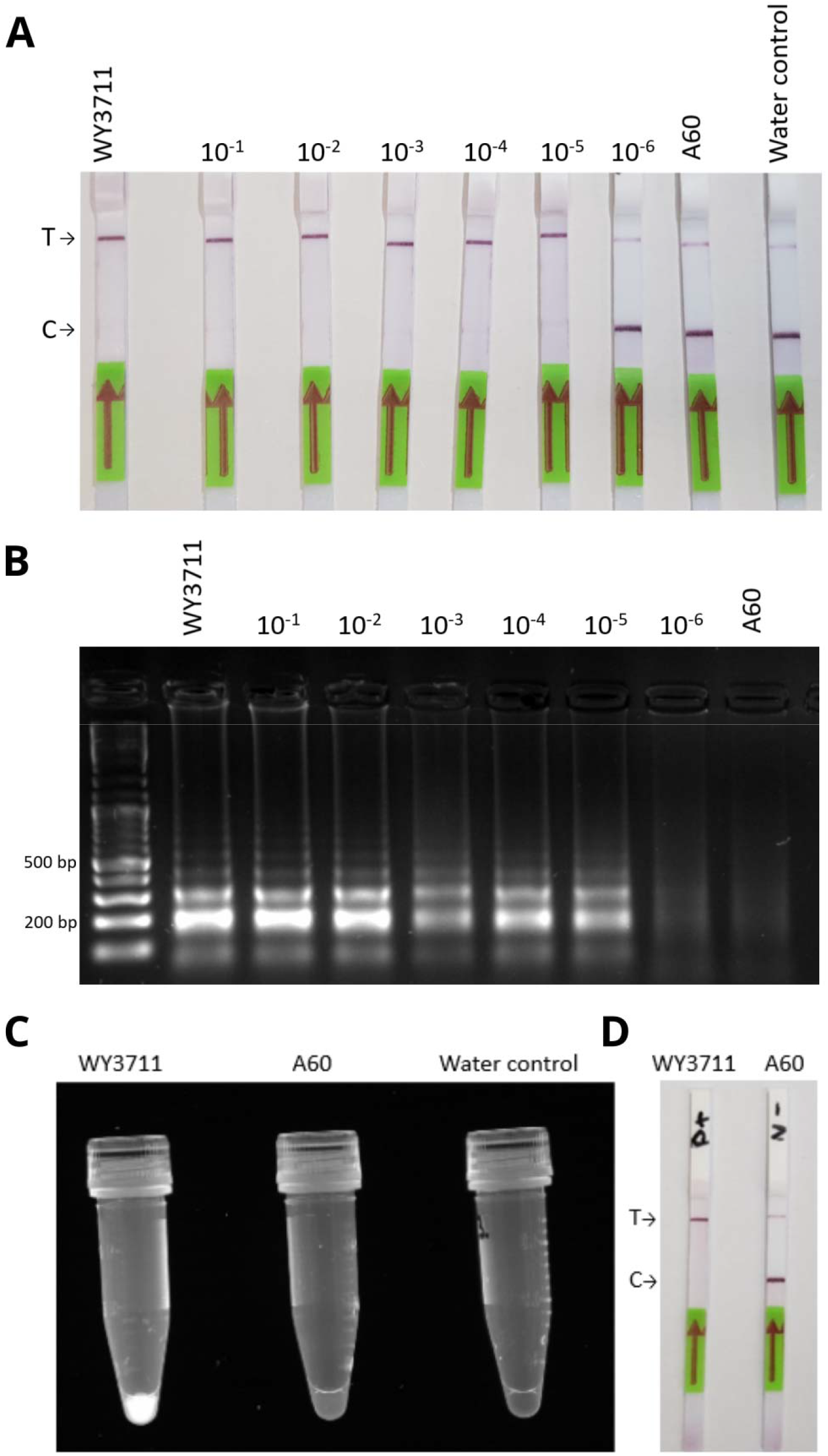
(**A)** Lateral flow read-out from the Cas12a reactions using DNA pre-amplified with LAMP from the dilution series (10^−1^ to 10^−6^) of *STA1+ S. cerevisiae* WY3711 in the *STA1*-*S. cerevisiae* A60. A NTC control was included where the DNA template was replaced with water. The T and C lines are marked next to lateral flow strips. (**B)** The corresponding LAMP products separated with gel electrophoresis. **(C**) Example of the Cas12a and LAMP reactions performed in the same tube using the F.Q. reporter and photographed under UV light. (**D**) Example of the Cas12a and LAMP reactions performed in the same tube using the biotin-labelled reporter and visualized with lateral flow read-out.

#### 3.2.3 One-pot assay development

To minimize pipetting steps and reduce cross-contamination risks, we next attempted to combine the LAMP and CRISPR-Cas12a reactions in a single reaction vessel. The LAMP reaction was assembled in the bottom of an Eppendorf tube and sealed with mineral oil, while the CRISPR-Cas12a reaction mix was prepared in the cap. Reactions were tested with positive (WY3711) and negative (A60) controls using both the F.Q. and biotin-labelled reporters. LAMP reactions were first incubated for 30 minutes at 65 °C, after which the Cas12a reaction mix was combined with the LAMP reaction inverting the tube and a brief centrifugation. The tubes were then incubated a further 30 minutes at 37 °C. The diastatic *S. cerevisiae* and non-diastatic *S. cerevisiae* were successfully distinguished with ‘one-pot’-method using both reporters (Figure 6C-D). With the fluorescent reporter, the positive sample was fluorescent under UV light (Figure 6C), while with the biotin-labelled reporter, the positive control produced only a strong T line on the lateral flow strips as observed previously (Figure 6D). A full protocol of the assay is available in Supplementary Note 1 in the Supplementary material.

#### 3.2.4 Discussion of the CRISPR-Cas12a assay

Results here showed that pre-amplification of template DNA was required to enable *STA1* detection through the Cas12a system. Out of the three tested methods, isothermal LAMP amplification was chosen as the most promising based on sensitivity. Ultimately, a successfully amplified *STA1* gene could be detected with a CRISPR-Cas12a reaction combined with two different visual detection methods (fluorescence and lateral flow). The *STA1* gene could also be detected directly using only LAMP amplification and gel electrophoresis as in Figure 4, and this has also been previously demonstrated (Hayashi et al. 2009), or as change in fluorescence or colour with DNA-binding dyes (Dao Thi et al. 2020). However, isothermal methods tend to have a high rate of non-specific amplification, but CRISPR-Cas systems, on the other hand, have high specificity and sensitivity (Mahas et al. 2021). LAMP was combined with CRISPR-Cas12a to improve reliability and accuracy.

### 3.3 Detection of diastatic *S. cerevisiae* from wort samples

To test whether the developed assay is applicable to wort or beer samples, we carried out wort fermentations that were intentionally contaminated with a diastatic strain. Three different wort fermentations were carried out in 12 °P wort in duplicate, and these were inoculated with either 1.2·10^7^ cells/mL of diastatic *S. cerevisiae* TPB109, non-diastatic *S. cerevisiae* WLP023, or WLP023 contaminated with 120 cells/mL of TPB109 (10^−5^ contamination). Based on sugar consumption and ethanol formation during fermentation, the TPB109 fermentation could be clearly differentiated from the WLP023 fermentation (Figure 7). This was particularly evident from the wort glucose concentrations, as glucose was rapidly consumed by WLP023, but present in low amounts almost throughout fermentation by the diastatic TPB109 due to glucoamylolytic activity. The contaminated fermentation could not be distinguished from the WLP023 fermentation from sugar and ethanol profiles alone. Presence of glucose in the contaminated fermentation could also not be observed.

**Figure 7.**
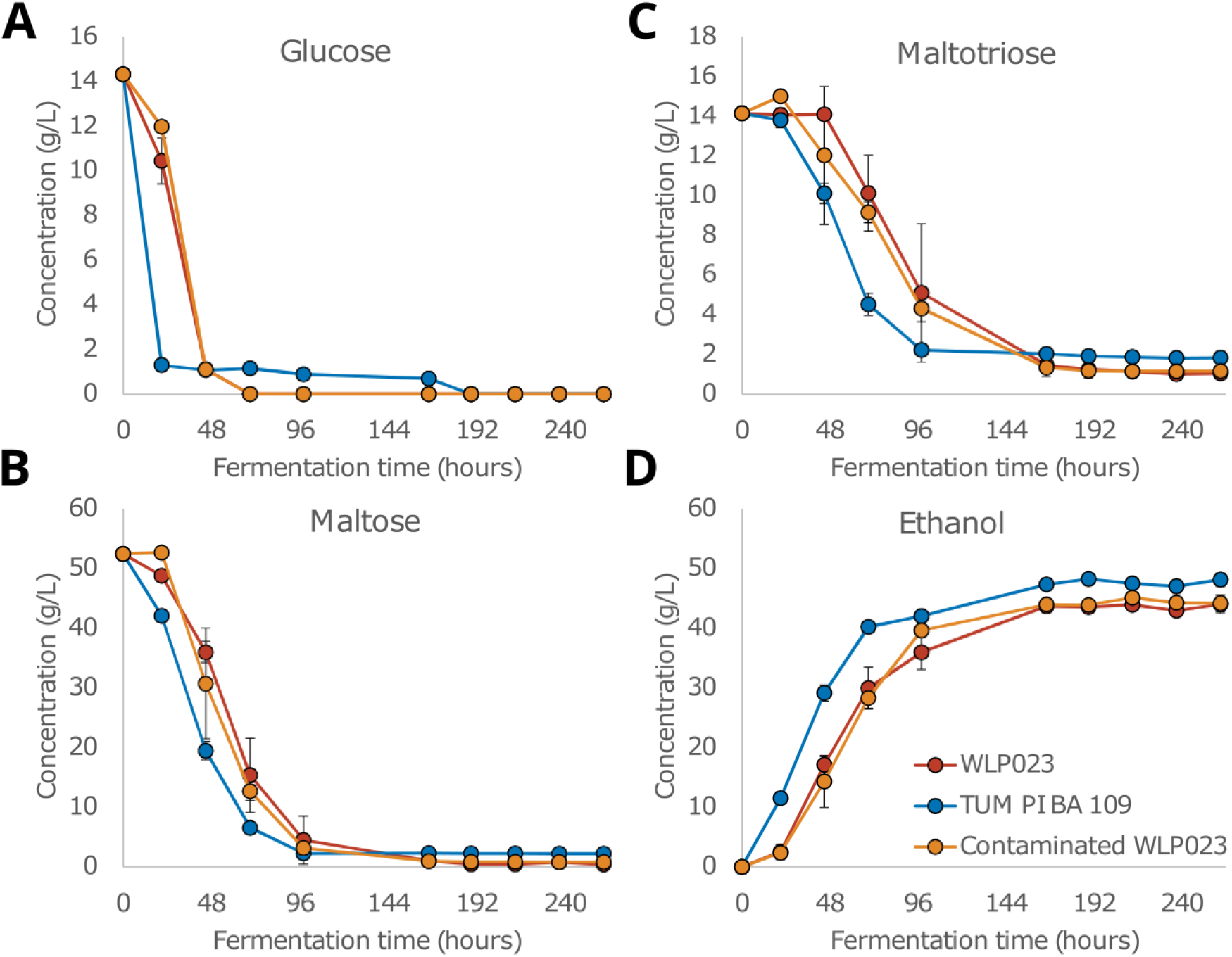
The concentrations (g/L) of (**A**) glucose, (**B**) maltose, (**C**) maltotriose, and (**D**) ethanol during fermentations in 12 °P wort. Fermentations were inoculated with *STA1-S. cerevisiae* WLP023 (red), *STA1+ S. cerevisiae* TUM PI BA 109 (blue), or WLP023 intentionally contaminated with TUM PI BA 109 (orange; contamination level of 10^−5^). Values are an average from two replicate fermentations and error bars represent the standard deviation.

DNA was extracted with ‘GC Prep’ from selected time points during fermentation. The selected time points were chosen from the beginning (2^nd^ day), middle (4^th^ and 7^th^ day), and end of the fermentation (11^th^ day). The selected samples included one of the two duplicate positive controls (TPB109), both duplicates of the contaminated fermentations (C1 and C2) and a negative control WLP023 from the last fermentation day. Sampling on 7^th^ day failed from one of the duplicate contaminated fermentations (C1), and the 7^th^ day C1 sample was therefore replaced with the corresponding 8^th^ day C1 sample. Extracted DNA samples were pre-amplified with both PCR and LAMP. The PCR results acted as a control for the LAMP-Cas12a assay, and both the commercially used SD5A/SD6B (Yamauchi et al. 1998) and the newly designed F2/R2 PCR primers were used.

With the SD5A/SD6B primers, contamination could only be detected from day 7 onwards (Figure 8A). With the F2/R2 primers, contamination could be detected already in the day 4 samples (Figure 8B). With LAMP, a product could also be detected with gel electrophoresis in the day 4 samples onwards (Figure 8C). However, bands were weak in some samples and difficult to distinguish from the negative control. After incubating the LAMP products with Cas12a, and visualizing results on lateral flow strips, clear positive results were again obtained for all contaminated samples from day 4 onwards (Figure 8D). Contaminated samples from day 2 also had slightly more intense T lines than the negative controls, indicating the potential for increased sensitivity of the assay compared to PCR, however, as the T-line could not be completely diminished from the negative controls, the C1 and C2 samples from the 2^nd^ day cannot be reliably interpreted as positive. By further optimizing and purifying the reporter oligo, to completely diminish the ‘hook effect’, or using different lateral flow strips, the assay sensitivity could be further improved.

**Figure 8.**
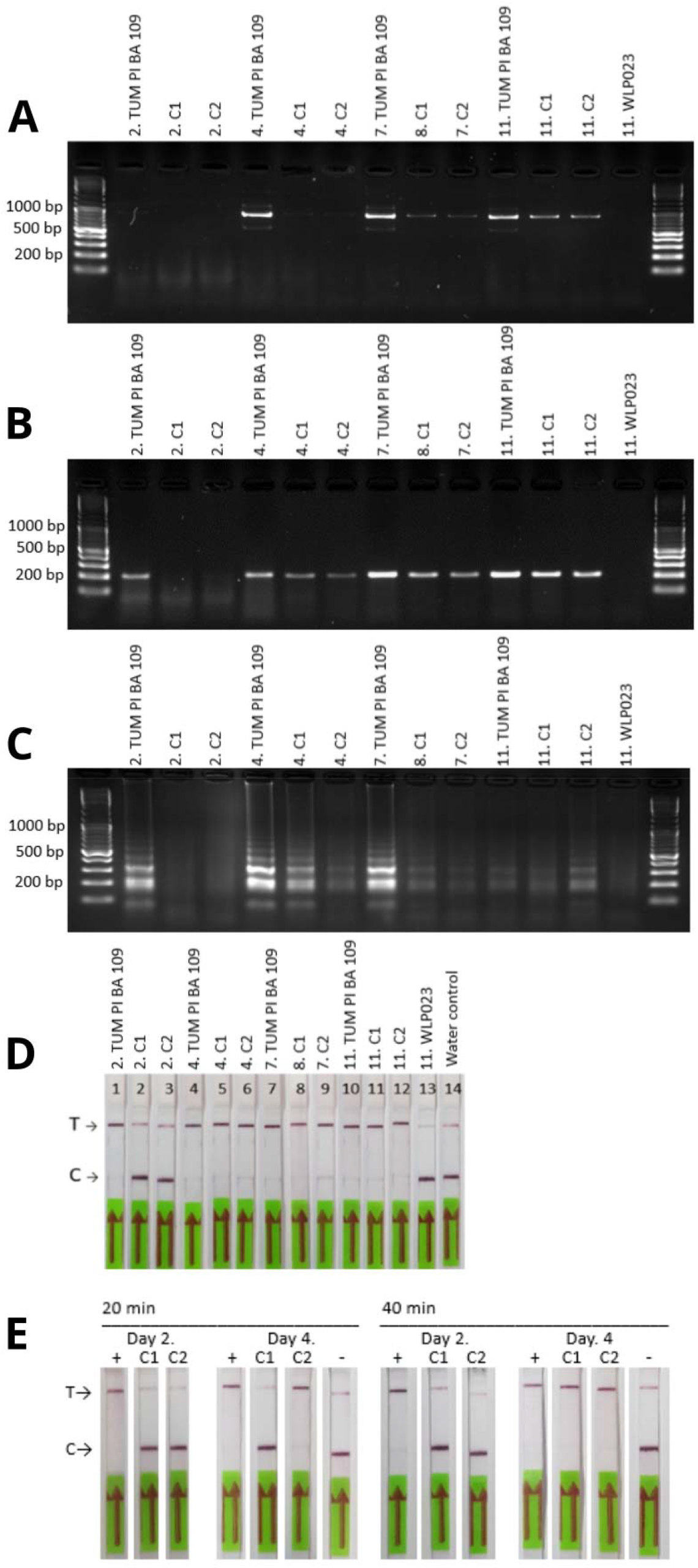
Detection of *STA1* from wort fermentations carried out with the *STA1*+ *S. cerevisiae* TPB109, *STA1-S. cerevisiae* WLP023, and WLP023 contaminated with TPB109 (C1 and C2). Numbers 2, 4, 7, 8 and 11 before sample name represent the day of fermentation when samples were collected. (**A**) Detection with PCR using SD5A/SD6B primers. (**B**) Detection with PCR using F2/R2 primers. (**C**) Detection with LAMP. (**D**) Detection with Cas12a reactions with lateral flow read-out on LAMP-pre-amplified DNA. (**E**) Detection with Cas12a reactions with lateral flow read-out on LAMP-pre-amplified DNA using altered incubation time (20 and 40 minutes) for the LAMP reaction.

The DNA samples from the second and fourth day of fermentation were retested with the one-pot method with lateral flow read-out. To also test whether assay results could be obtained more quickly or with greater sensitivity, LAMP incubation time was decreased to 20 and increased to 40 minutes, respectively. Results indicate that a shorter 20 minute incubation time during LAMP is sufficient for a positive result in samples with high amounts of diastatic yeast, such as the positive controls (Figure 8E). However, only one of the 4^th^ day contaminated samples C2 was detected as positive with the shorter incubation time. Results with the 40 minute incubation time during LAMP were identical to those obtained with 30 minute incubation (data not shown), and therefore sensitivity could not be improved by increasing the LAMP incubation time.

## 4. Conclusions

Diastatic *S. cerevisiae* is one of the most common contaminating microbes in beer fermentations, especially in smaller breweries with less stringent quality control and lack of pasteurization equipment. To ensure high beer quality, it is therefore vital to enable rapid and accurate detection of these yeasts in brewery samples; preferably without the need of expensive equipment. The objective of this study was to develop such an easy and rapid detection assay for diastatic *S. cerevisiae* using only basic laboratory equipment that are available in most laboratories. Such an assay be easily implemented for quality control in both microbreweries and industrial-scale breweries.

Here, we successfully developed a rapid detection assay for diastatic *S. cerevisiae* using nucleic acid detection by CRISPR-Cas12a. The developed assay consisted of three steps: DNA extraction with ‘GC Prep’, isothermal pre-amplification with LAMP, and CRISPR-Cas12a-based nucleic acid detection with lateral flow readout. With this assay, visually detectable results were achieved within 75 minutes from yeast samples using only pipettes, pipette tips, tubes, a vortex mixer, a heat block, a lateral flow kit and reagents involved in DNA extraction and the LAMP-Cas12a assay. Compared to other molecular detection techniques, such as PCR or quantitative PCR, this new assay does not need, for example, a thermal cycler, gel electrophoresis devices, or a qPCR machine. While the new assay yields rapid results, we were unsuccessful in our aim to achieve results within 1 hour. However, we believe the reactions and their timings could be further optimised to further shorten the overall assay time.

The developed assay could detect contamination at levels of 10^−5^ with both fluorescence and lateral flow strips when yeast suspensions in water were prepared. The assay was also tested on wort fermentations intentionally contaminated with diastatic *S. cerevisiae*, and from an initial contamination level of 10^−5^, contamination could be detected from day 4. These detection limits were equivalent to those obtained using PCR, and are similar to reported limits in a previous study, where industrial brewing samples were spiked with diastatic *S. cerevisiae* and 0.001% of contamination was detected with real-time PCR (Michel et al. 2016). Curiously, when the samples from the wort fermentations were analysed, detection was more reliable with the new assay compared to the detection by PCR using the widely used SD-5A/SD-6B primers. Overall, this new assay can be applied for rapid detection of diastatic yeast in any brewery, but we believe this new assay will be particularly useful for smaller breweries that don’t already have well-equipped laboratories and are looking to implement better quality control.

## Author Contributions

IU: designed experiments, performed experiments, analysed the data, and wrote the manuscript

KK: conceived the study, acquired funding, designed experiments, analysed the data, and wrote the manuscript.

## Acknowledgements

IU was funded by Tor-Magnus Enari Foundation, and the work was performed as a part of her MSc thesis.

