## Supplementary Material for "A simple and rapid CRISPR-Cas12a-based detection test for diastatic *Saccharomyces cerevisiae*"

**Supplementary Table 1.** Primers designed to RPA reactions.

| **Name** | **Sequence (5′ > 3′)** |
| --- | --- |
| STA1_RPA_F1 | AGTTGTTGAAGGGTTCTCAATTGATAAAAAAG |
| STA1_RPA_F2 | TTACCTGTACAAGTTGTTGAAGGGTTCTCAA |
| STA1_RPA_F3 | CCTGTACAAGTTGTTGAAGGGTTCTCAATTGATAAAAAAG |
| STA1_RPA_R1 | GAAGTGATGTTGCTAGAGGAGGATCCTCTAGGAACT |
| STA1_RPA_R2 | GAAGTGATGTTGCTAGAGGAGGATCCTCTA |
| STA1_RPA_R3 | ATGTTGCTAGAGGAGGATCCTCTAGGAACT |
| STA1_RPA_R4 | TGTGCGTATATGGATTTTTGAGGCCTACCA |

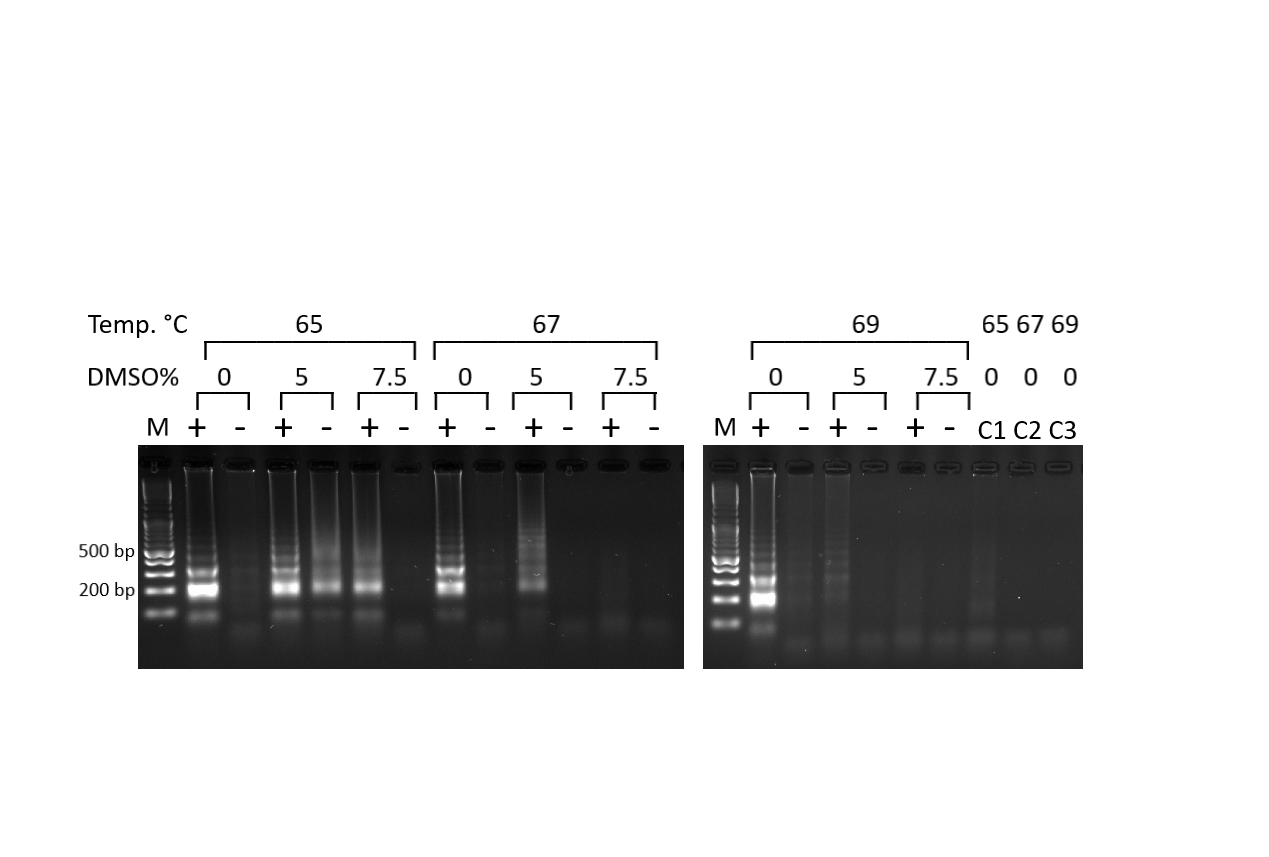

**Supplementary Figure 1.** Optimization of the LAMP reaction conditions with DMSO contents of 0%, 5%, 7.5% and temperatures 65 °C, 67 °C and 69 °C. Optimization was performed with WY3711 as the *STA1* positive control (+) and A60 as the negative control (-). The DNA ladder is marked as (M). C1, C2 and C2 are no-template controls where DNA is replaced with water.

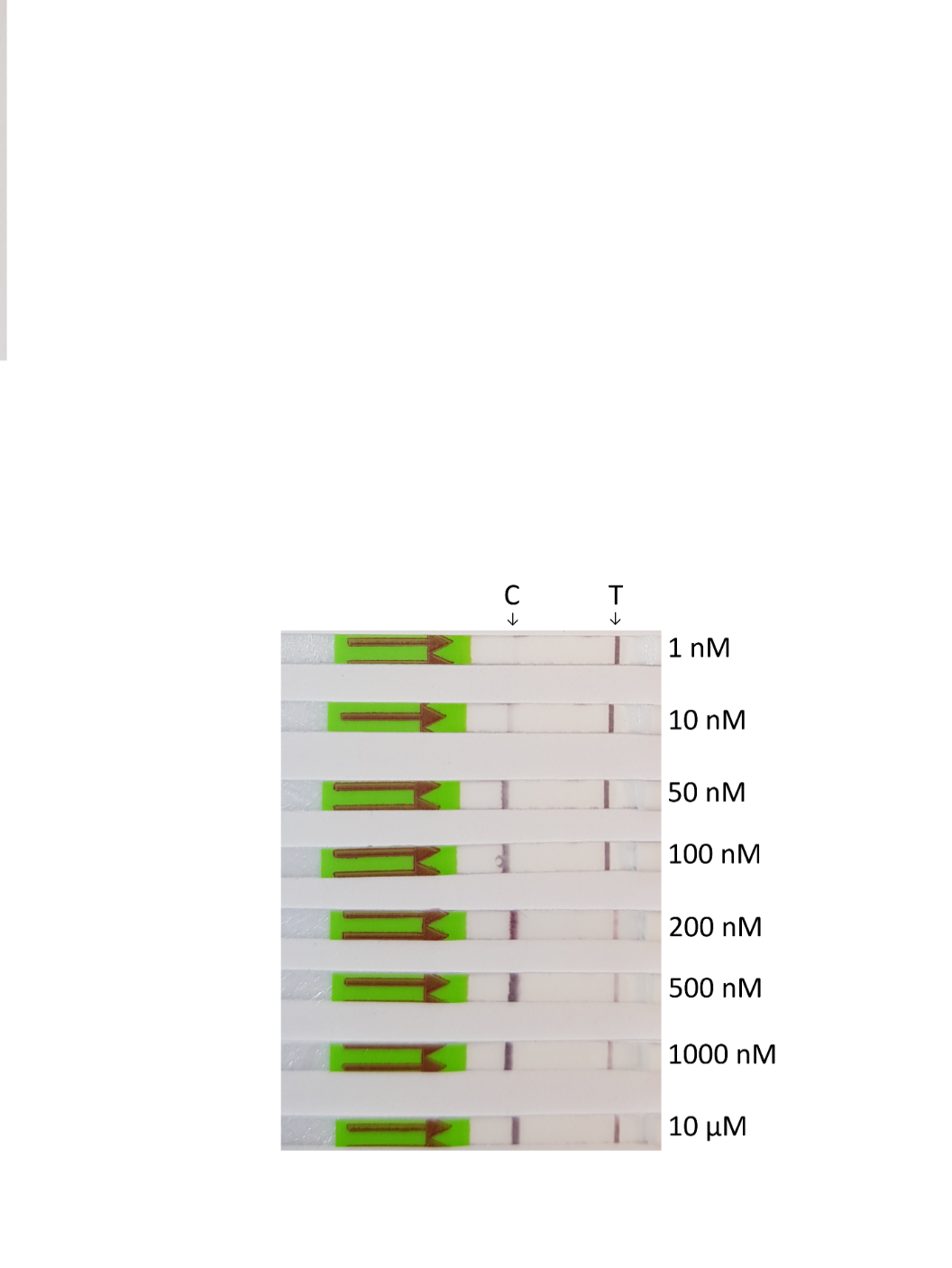

**Supplementary Figure 2.** Testing of the ‘hook effect’ by transferring aqueous solutions of the biotin-labelled reporter oligo at different concentrations to the lateral flow strip. The oligo was HPLC-purified.

**Supplementary note 1:**

A full protocol for the detection of *STA1* from a yeast/beer sample using LAMP and CRISPR-Cas12a.

***Step 1: DNA Extraction (10 minutes)***

***Materials:***
5% Chelex 100 in deionized water
Acid washed glass beads

1. Transfer 100 µL of yeast culture into a sterile 1.5mL Eppendorf tube.
2. Centrifuge for 1 minute at 5000 × *g*. Remove supernatant.
3. Resuspend pellet in 100 μl 5% Chelex 100, and add glass beads to half total sample volume (~100 µL).
4. Vortex mix at high speed for 4 minutes.
5. Incubate sample at 98 °C for 2 minutes.
6. Spin microtube at top speed (15000 × *g*) in a microcentrifuge for 1 minute.
7. Collect 70 µL of supernatant in a clean tube, ensuring that no Chelex 100 is carried over.

***Step 2: LAMP + Cas12a one-pot reaction (63 minutes)***

***Materials:***
New England Biolabs WarmStart LAMP 2X Master Mix

LAMP Primer Mix (10X concentration)

Nuclease-free water

NEBuffer r2.1 Reaction Buffer (10X)

crRNA (10 µM)

EnGen® Lba Cas12a (10 µM)

Biotin-labelled reporter oligo (5' / 6-FAM / TTATT / Biotin / 3'; 10 µM)

TaKaRa Bio RNase inhibitor

1. Prepare the LAMP reaction in a 1.5ml RNase-free tube (reaction volume 10 µl) as in the table below. 1 µl of template DNA is added to reaction and mixed. The reaction mix is covered with mineral oil (e.g. 25 µl).

|  | Per reaction |
| --- | --- |
| WarmStart LAMP 2X Master Mix | 5 µl |
| LAMP Primer Mix (10X) | 1 µl |
| Nuclease-free water | 3 µl |
| DNA extracted in Step 1 | 1 µl |

2. Assemble the Cas12a reaction mix in the lid of the same tube:

|  | Per reaction |
| --- | --- |
| Nuclease-free water | 12 µL |
| NEBuffer r2.1 Reaction Buffer (10X) | 3 µL |
| crRNA (10 µM) | 1.5 µL |
| Cas12a enzyme (10 µM) | 1.5 µL |
| Reporter oligo (10 µM) | 1.2 µL |
| RNase inhibitor | 0.75 µL |

3. Incubate the tube at 65 °C for 30 minutes. The tube is inverted five times, to mix the LAMP and Cas12a reaction mixes. The tube is quickly spun down in a centrifuge. The Cas12a reaction is then incubated at 37 °C for 30 minutes.
4. If testing multiple samples, LAMP and Cas12a reaction mixes can be prepared in larger volume and divided to the tubes.

***Step 3: Visualisation on a lateral flow strip (2 minutes)***

***Materials:***
HybriDetect - Universal Lateral Flow Assay Kit

1. A 5 µL sample from the LAMP/Cas12a reaction is transferred to the application area of lateral flow strip. The strips are incubated for 1 min in 100 µL of HybriDetect assay buffer in an upright position and results were observed immediately or photographed.
